# The Effect of Gradual Ovarian Failure on Dynamic Muscle Function and the Role of High Intensity Interval Training on Mitigating Impairments

**DOI:** 10.1101/2023.07.14.548999

**Authors:** Emma F. Hubbard, Parastoo Mashouri, W Glen Pyle, Geoffrey A Power

## Abstract

Skeletal muscle function is impaired in menopause and exercise may mitigate tshis decline. We used the VCD model of menopause to investigate the effects of gradual ovarian failure on skeletal muscle contractile function and whether high intensity interval training (HIIT) can mitigate impairments. Sexually mature female CD-1 mice were assigned to one of three groups: control (n=5), VCD-sedentary (n=5), or VCD-training (n=5). Following ovarian failure, the VCD-training group underwent 8 weeks of uphill HIIT. Mice were sacrificed 8 weeks after ovarian failure, representing late menopause. Single muscle fibres from the soleus (SOL) and extensor digitorum longus (EDL) muscles were dissected, chemically permeabilized, and mechanically tested. Single muscle fibres were maximally activated (pCa 4.5) then isotonic load clamps were performed to calculate force-velocity-power curves. Absolute force and peak power were 31% and 32% lower in VCD-sedentary fibres compared to control fibres, respectively, in both SOL and EDL muscles. Despite reductions in absolute force and therefore lighter relative loads imposed during the isotonic contractions in VCD-sedentary fibres, there were no concomitant increases in contractile velocity. HIIT was partially effective at mitigating power loss (22% higher peak power in VCD-training compared to VCD-sedentary), but only in fast-type SOL fibres. These findings indicate that ovarian failure impairs dynamic contractile function – likely through a combination of lower force-generating capacity and slower shortening velocity – and that HIIT may be insufficient to completely counteract the deleterious effects of menopause at the cellular level.

**New & Noteworthy:**

- Reductions in circulating ovarian hormones impair static muscle contractile performance, but less is known about dynamic properties like power.
- Typically, rodent models of menopause completely remove the ovaries and fail to mimic the prolonged and complex hormonal transition that includes a retention in ovarian androgen production.
- Using an ovary intact VCD model of ovarian failure, we found that single fibre power was impaired compared with controls in both SOL and EDL fibres.
- Our uphill high intensity interval training program was partially sufficient to reverse power loss, but only in fast-type SOL fibres.
- Impairments in muscle power following ovarian failure are likely driven by a combination of decreased muscle size and force-generating capacity.

## Introduction

The loss of ovarian reproductive function, termed menopause, occurs with ovarian follicular exhaustion, which results in a rapid and dramatic decrease in the synthesis of 17β-estradiol (1, 2). The ovariectomized (OVX) rodent model is commonly used to assess the effects of reduced circulating hormone levels on muscle function; however, this model is limited by the complete excision of all ovarian tissue, which does not occur in the natural progression of menopause, and the immediate hormonal shift which fails to mimic the transitional phase known as perimenopause (3). The occupational chemical 4-vinylcyclohexene diepoxide (VCD) provides an alternative model of menopause by accelerating the natural process of atresia, thereby producing a follicle-deplete, ovary-intact animal that includes a retention in androgen production (3, 4). Recent work from our lab showed that immediately after VCD-induced ovarian failure (i.e., onset of menopause) in young female mice, muscle function was not significantly impaired compared to controls (5), thus demonstrating the importance of evaluating the effects of VCD-induced ovarian failure on muscle contractility beyond the onset of follicular exhaustion.

Isometric force production has been shown to be lower in both single muscle fibres (6; 10-14 weeks post-surgery) and whole muscle preparations (7; 8-9 weeks post-surgery) from OVX rats and mice, respectively, compared to their ovary-intact counterparts. Across studies, normalized muscle force (indicative of muscle quality), was ∼7% greater in estrogen-replaced mice compared to OVX mice (8). While recent work from our lab did not observe any reductions in force immediately following VCD-induced ovarian failure (i.e., onset of menopause) (5), Greising et al. (9) found that 8 weeks after ovarian failure estrogen-replaced mice produced 16-19% more concentric, isometric, and eccentric forces compared to non-replaced VCD mice, indicating that circulating 17β-estradiol plays a key role in muscle contractile function independent of the presence of ovarian tissue. It is commonly thought that hormone-related reductions in force production capacity are related to decreases in skeletal muscle myosin regulatory light chain phosphorylation following OVX which reduces the probability of cross-bridge attachment (10, 11). Additionally, estrogens have been shown to have membrane stabilizing properties that prevent damage to the muscle membrane caused by exercise (12) or oxidative stress (13), which may provide an explanation for the detrimental effect of hormone reduction on muscle quality.

Muscular power, which is the product of both force and velocity, is a better predictor of functional mobility than independent measures of force or velocity (14). At the whole muscle level, power production across muscle groups has been shown to be unchanged following OVX (7; 8-9 weeks post-surgery) and impaired in skeletal muscle estrogen receptor knockout models (15) as compared to controls. Interestingly, the largest impairments in power seem to occur at slower shortening velocities (15), indicating that force loss may be driving some of the observed impairments in power more so than velocity. In fact, faster contractile velocity has been reported in the SOL muscle of OVX mice compared to controls, indicating that impairments in muscle power production could even be offset by increases in shortening velocity (7; 8-9 weeks post-surgery). However, it remains unclear how gradual ovarian failure and reduced levels of circulating ovarian hormones affect dynamic properties of muscle function, particularly at the cellular level.

Various modes of exercise training have been offered as countermeasures to attenuate reductions in muscle mass and quality associated with aging and hormonal changes (16). Long-term moderate-to-vigorous aerobic exercise has been shown to yield enhancements in single muscle fibre power production in menopausal females through fibre type-specific mechanisms, with type I (i.e. slow-twitch) fibre power being improved primarily though increases in force production and type II (i.e. fast-twitch) fibre power being maintained primarily through increases in contractile velocity (17). Following 14 weeks of uphill endurance running in young OVX rats, SOL and EDL muscle mass and twitch contraction speed increased, as did SOL tetanic force (18; 14 weeks post-surgery). Uphill high intensity interval training (HIIT) protocols have been successful in slowing atrophy and functional impairments in aged male and female mice with smaller reductions in SOL and EDL muscle size and force production compared to age-matched control animals, as well as improvements in gait speed, endurance, and anaerobic capacity (19, 20). As such, it stands to reason that a similar uphill HIIT protocol may mitigate any impairments in muscle function owing to a gradual reduction of circulating ovarian hormones.

The purpose of the present study was to investigate the effects of VCD-induced gradual ovarian failure on single muscle fibre power production in late-stage menopause and identify fibre phenotype-specific effects. Additionally, this study aimed to determine whether an uphill HIIT running protocol could slow or reverse impairments in dynamic muscle contractile function owing to ovarian failure. It was hypothesized that single muscle fibre power would be lower in the VCD-sedentary group compared to the sedentary control group, and that HIIT would mitigate these changes.

## Materials & Methods

### Animals

Fifteen sexually mature female CD-1 mice (sacrificial age ∼ 34 weeks; mass = 44.1 ± 7.9 g) were obtained (Charles River Laboratories, Senneville, QC, Canada) with approval from the University of Guelph’s Animal Care Committee and all protocols followed CCAC guidelines. Mice were housed at 24°C in groups of four and given ad-libitum access to food and room-temperature water. A 12-hour light/12-hour dark cycle was used, and no technicians entered the room during the dark cycle. Following one week of acclimation to housing conditions, mice were assigned to one of three experimental groups: control (n=5), VCD-sedentary (n=5) and VCD-training (n=5). Mice in both VCD groups were injected intraperitoneally with 160 mg/kg body weight of VCD (diluted with sesame oil to a 0.0587:1 ratio) daily for 15 days. 110 days after the first injection, vaginal swabs were performed to ensure that mice remained in the diestrus phase for ten consecutive days, indicating that ovarian failure did in fact occur. Sacrifice by CO_2_ asphyxiation occurred 8 weeks after ovarian failure, representing a late menopause timepoint.

### Training Protocol

Two weeks prior to training, mice in the VCD-training group began treadmill familiarization. During the first week of familiarization, the mice ran on a flat EXER 3/6 animal treadmill (Columbus Instruments) on three different days for 5 minutes, beginning at a speed of 5 m/min and accelerating 1 m/min to a 10 m/min finish. During the second week, the mice repeated the same familiarization protocol at a 25° incline. Training was initiated 120 days after VCD injection, following confirmation of ovarian failure. The training protocol consisted of eight weeks of 10-minute uphill high intensity interval training sessions at a 25° incline, 3 days/week. The training protocol employed is adapted from the training intervention outlined by Seldeen et al (19). Briefly, each training session consisted of a 3-minute warm-up at a base speed that was easily maintained by all mice, then three 1-minute intervals at a sprint speed (with 1 minute of recovery at base speed between sprints) and a final fastest 1-minute dash interval. Intensity was increased every two weeks and all speeds were adjusted accordingly (Figure 1).

**Figure 1:**
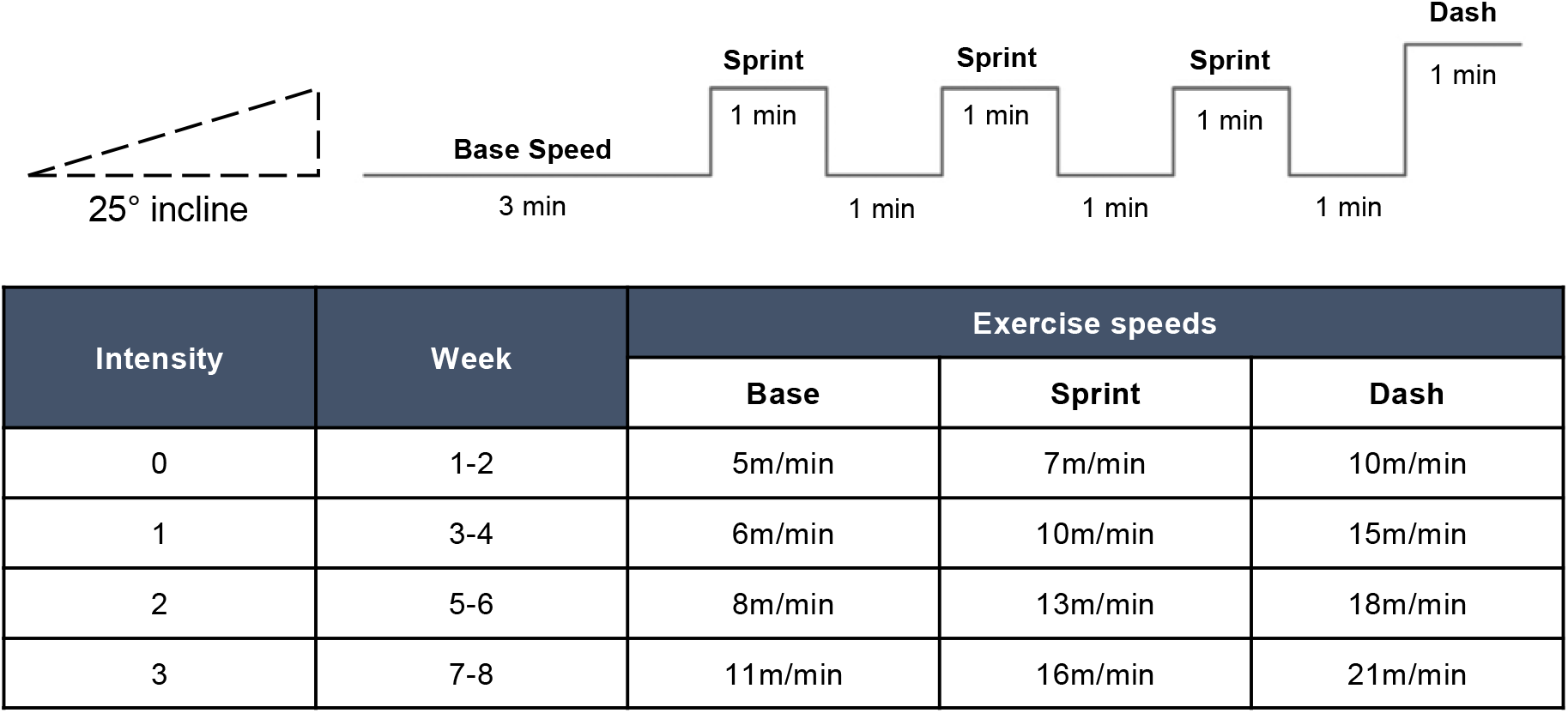
High intensity interval training program and intensity progression.

### Muscle Preparation

Soleus (SOL) and extensor digitorum longus (EDL) muscles were harvested from mice following sacrifice 8 weeks after ovarian failure, representing a timepoint associated with late menopause. Muscles were submerged in chilled dissecting solution. These muscles were selected to represent distinct fibre types as the mouse SOL muscle is composed primarily of type I muscle fibres (75%) and the EDL is almost entirely type II muscle fibres (>90%), allowing any phenotypic dependency to be observed (21, 22). Additionally, both mechanical and metabolic adaptations have been observed in mouse SOL and EDL muscles following treadmill running and high intensity interval training protocols (23). While in the dissecting solution, several muscle fibre bundles of approximately 0.5–1 mm in width and 3 mm in length were dissected and transferred to a tube containing 2.5 mL of chilled skinning solution on ice for 30 minutes to be chemically permeabilized (24). Gentle agitation was performed to ensure that all bundles were constantly submerged in the solution, resulting in equal permeabilization. The bundles were then washed with fresh chilled dissecting solution, gently agitated to remove any remaining skinning solution, and stored in a separate tube containing fresh storage solution. The bundles were incubated for 24 hours at 4°C. Tubes were prepared with fresh storage solution and bundles were placed separately in each tube and placed in a freezer at −80°C until mechanical testing (25).

### Solutions

The dissecting solution was composed of the following (in mM): K-proprionate (250), Imidazole (40), EGTA (10), MgCl_2_•6H_2_O (4), and Na_2_H_2_ATP (2). The storage solution was composed of K-propionate (250), Imidazole (40), EGTA (10), MgCl_2_•6H_2_O (4), Na_2_H_2_ATP (2), glycerol (50% of total volume after transfer to 50:50 dissecting:glycerol solution). The skinning solution with Brij 58 was composed of K-propionate (250), Imidazole (40), EGTA (10), MgCl_2_•6H_2_O (4), 1 g of Brij 58 (0.5% w/v). The relaxing solution was composed of Imidazole (59.4), KMSA (86), Ca(MSA)_2_ (0.13), Mg(MSA)_2_ (10.8), K_3_EGTA (5.5), KH_2_PO_4_ (1), Leupeptin (0.05), and Na_2_ATP (5.1). The pre-activating solution consisted of K-propionate (185), MOPS (20), Mg(CH_3_COO)_2_ (2.5), and Na_2_ATP (2.5). The activating solution (pCa 4.5) consisted of Ca^2+^ (15.11), Mg (6.93), EGTA (15), MOPS (80), Na_2_ATP (5), and CP (15). All solutions were adjusted to a pH of 7.0 with the appropriate acid (HCl) or base (KOH).

### Mechanical Testing

On the day of mechanical testing, muscle bundles were removed from the freezer and placed in a chilled relaxing solution on ice where they remained for the duration of the testing protocol. Single fibres were dissected from the bundles in relaxing solution, then transferred into a temperature-controlled chamber (15°C) filled with relaxing solution, and tied with nylon suture knots between a force transducer (model 403A; Aurora Scientific) and a length controller (model 322C; Aurora Scientific). The average sarcomere length (SL) was measured using a high-speed camera (Aurora Scientific). Before starting the testing protocol, a ‘fitness’ contraction was performed at a SL of 2.5 μm (26) in pCa 4.5 to ensure the ties were not loose and the fibre did not have extreme SL non-uniformity caused by damage. After the fitness test, SL was re-measured and, if necessary, re-adjusted to 2.5μm. Fibre length (L_0_) was recorded, and fibre diameter was measured at three different points along the fibre using a reticule on the microscope. The average of the fibre diameters was used to calculate cross-sectional area (CSA) assuming circularity. To measure maximal isometric force production (P_o_), fibres were transferred to a pre-activating solution (reduced Ca^2+^ buffering capacity with ATP) for 20 seconds, then to an activating solution (pCa 4.5) for 25 seconds. P_o_ was recorded and used to determine submaximal isotonic load clamp values. Instantaneous stiffness was determined as a change in force for a given change in length, calculated as (-) 0.5% L_0_. Fibres were maximally activated in pCa 4.5 before applying each isotonic protocol. Isotonic load clamps were performed through 2 activations of the fibre (separated by 60 seconds of rest between activations) with 4 load clamps each for a total of 8 loads (10, 20, 30, 40, 50, 60, 70, and 80% P_o_). Isotonic load clamps were performed in a random order but always from highest to lowest force within a single activation.

### Analysis

To determine the force-velocity relationship, isotonic load clamps were performed at 8 submaximal force levels (10, 20, 30, 40, 50, 60, 70, and 80% P_o_) and the resultant length changes were measured over 0.45 seconds. Shortening velocity for each % P_o_ was calculated as ΔL/Δt and data across all loads was fit to a force-velocity curve using a modified hyperbolic Hill equation (27):

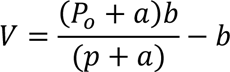

where *P* is load, *V* is velocity, *P_0_* is the isometric force, and *a* and *b* are constants (Figure 2). All force values were normalized to CSA. The force-velocity data was modeled to the hyperbolic Hill equation (27) such that the sum of the squares of the deviations of predicted velocity from observed velocity was minimized. Goodness of fit was determined by the R-squared (R^2^) statistic. A threshold goodness of fit value was set at R^2^ = 0.90 and any curves with values below this were removed as outliers. Based on this criteria, *n*= 20 control fibres, *n* = 27 VCD-sedentary fibres, and *n* = 17 VCD-training fibres were excluded. The ratio *a*/*P*_0_ indicates the curvature of the force-velocity relationship. Maximal shortening velocity, V_max_ was extrapolated from the fit force-velocity curve where load was equal to 0. Absolute power was defined as the product of force (in mN/mm^2^) and shortening velocity (in L_0_/s) yielding final units of mN•mm^-2^•L_0_•s^-1^ and a force-power curve was also constructed. Peak power was calculated from the force-power relationship to compare single muscle fibre power between groups. For each outcome measure, any value beyond 3 standard deviations from the mean was removed as an outlier.

**Figure 2:**
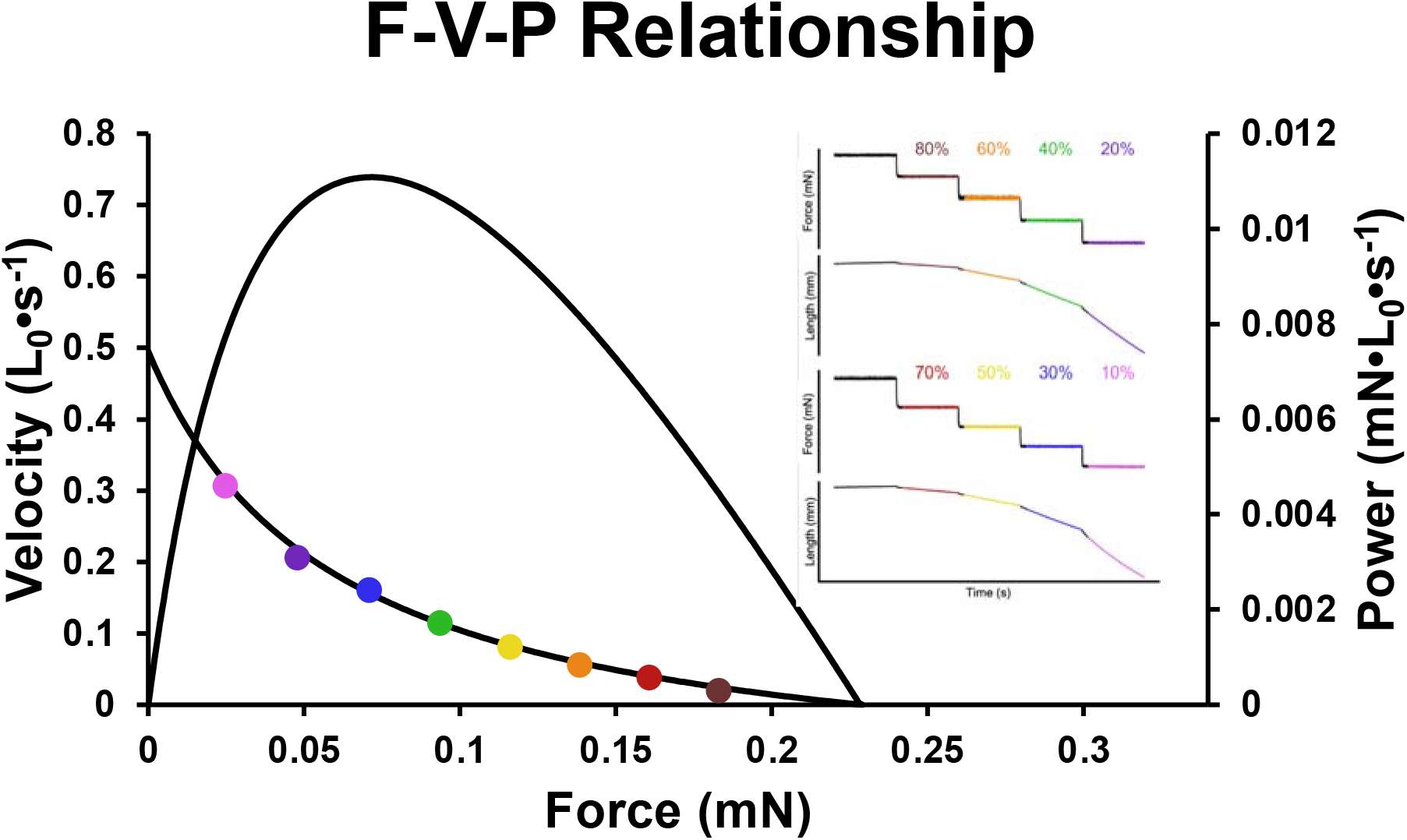
Example force-velocity-power curve from submaximal isotonic load clamps. Length and force data obtained during a series of isotonic contractions over two maximal Ca^2+^ activations. The fibre had attained peak isometric force (P_0_) at beginning of trace and was then allowed to shorten – as indicated by the downward deflection of the length record – to achieve a set % of P_0_. Fibre force and shortening velocity were determined over the final 450 ms of each step, corresponding to the linear portion of the length change.

### Fibre Type

SOL muscles are primarily composed of MHC type I ‘slow’ fibres (∼75%) while EDL muscles are almost entirely MHC type II ‘fast’ fibres (>90%) (21, 22). However, there appears to be a disconnect between MHC biochemical fibre type and mechanical phenotype with a multitude of genes and transcription factors implicated in mechanical phenotypic regulation (28, 29). As such, fibres were grouped by mechanical phenotype rather than MHC isoforms. “Binning” of muscle fibres has been performed on human fibres based on unloaded shortening velocity (V_0_) (30, 31) and V_max_ has been shown to be closely related to MHC fibre type in skinned rat fibres (32). In a post-hoc analysis, muscle fibres were phenotypically binned into a ‘slow’ or ‘fast’ type group based on V_max_ values obtained in each of the three groups (*n* fibres for: control; 78, VCD-sedentary; 72, VCD-training; 81). Fibres slower than 40% of the fastest V_max_ in each group (after removing outliers) were considered ‘slow’ and binned into the slow-type fibre group, and fibres faster than 40% of the fastest V_max_ in each group (after removing outliers) were considered ‘fast’ and binned into the fast-type fibre group.

### Statistical Analysis

For all statistical analysis, single muscle fibres were treated as independent samples, with 10 fibres sampled from each muscle from each mouse yielding a total sample size of *n* = 300 fibres. Muscle contractility (force, power) and size (CSA) were assessed using a two-way ANOVA with group and muscle as between-group factors. To identify any fibre-type differences, a secondary two-way ANOVA was performed with group and fibre phenotype as between-group factors. Significance was set at an alpha of 0.05 and all analyses were completed with GraphPad Prism (GraphPad Software, LLC.; Version 9.2). All data in figures are presented as mean ± SEM.

## Results

### Absolute force

For absolute force, there was no interaction of group × muscle (*p* = 0.1192), but there was a main effect of group (*p* < 0.0001) such that VCD-sedentary fibres produced 31.0% less force compared to control fibres and 24.2% less force compared to VCD-training fibres (Figure 3A). There was no main effect of muscle (*p* = 0.4054). For the SOL, when fibres were binned based on V_max_, there was no interaction of group × phenotype (*p* = 0.0957), but there was a main effect of group (*p* = 0.0003) such than VCD-sedentary SOL fibres produced 26.7% less force than control SOL fibres (Figure 3B). There was also a main effect of phenotype (*p* < 0.0001) such that fast SOL fibres produced 29.0% more force than slow SOL fibres. For the EDL, when fibres were binned based on V_max_, there was no interaction of group × phenotype (*p* = 0.18250), but there was a main effect of group (*p* = 0.0140) such that VCD-sedentary EDL fibres produced 32.9% less force than control EDL fibres (Figure 3C). There was no effect of phenotype (*p* = 0.5191).

**Figure 3:**
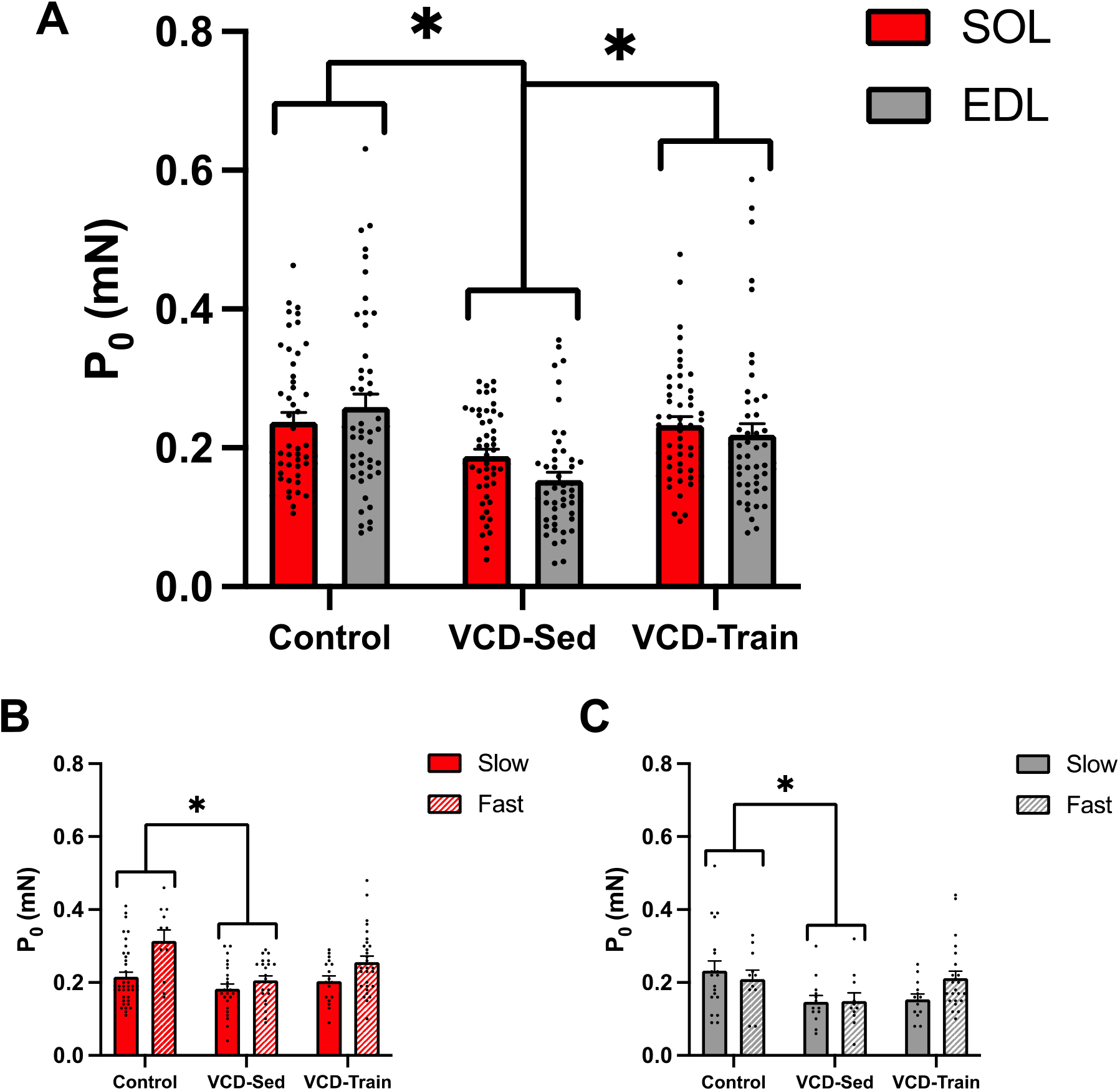
A) Absolute force. Comparison of peak isometric force (P_0_, mN) from maximally activated (pCa 4.5) single SOL (red) and EDL (grey) muscle fibres. VCD-sedentary fibres produced less absolute force compared to control and VCD-training fibres. **B) SOL muscle.** When analyzing the SOL muscle by fibre phenotype, VCD-sedentary fibres produced less absolute force than controls, and fast fibres (hashed) produced more absolute force than slow fibres (solid). **C) EDL muscle.** When analyzing the EDL muscle by fibre phenotype, VCD-sedentary fibres produced less absolute force than controls and there was no effect of phenotype. **\*** indicates significant difference (*p* < 0.05).

### Cross-sectional area

For CSA, there was no interaction of group × muscle (*p* = 0.0634), but there was a main effect of group (*p* = 0.0003) such that VCD-sedentary fibres were 21.8% smaller than control fibres and 14.1% smaller than VCD-training fibres (Figure 4A). There was also a main effect of muscle (*p* = 0.0005) such that EDL fibres were 18.8% larger than SOL fibres. For the SOL, when fibres were binned based on V_max_, there was no interaction of group × phenotype (*p* = 0.4282), nor main effects of group (*p* = 0.1670) or phenotype (*p* = 0.2601) (Figure 4B). For the EDL, when fibres were binned based on V_max_, there was no interaction of group × phenotype (*p* = 0.4174), nor main effects of group (*p* = 0.0774) or phenotype (*p* = 0.6012) (Figure 4C).

**Figure 4:**
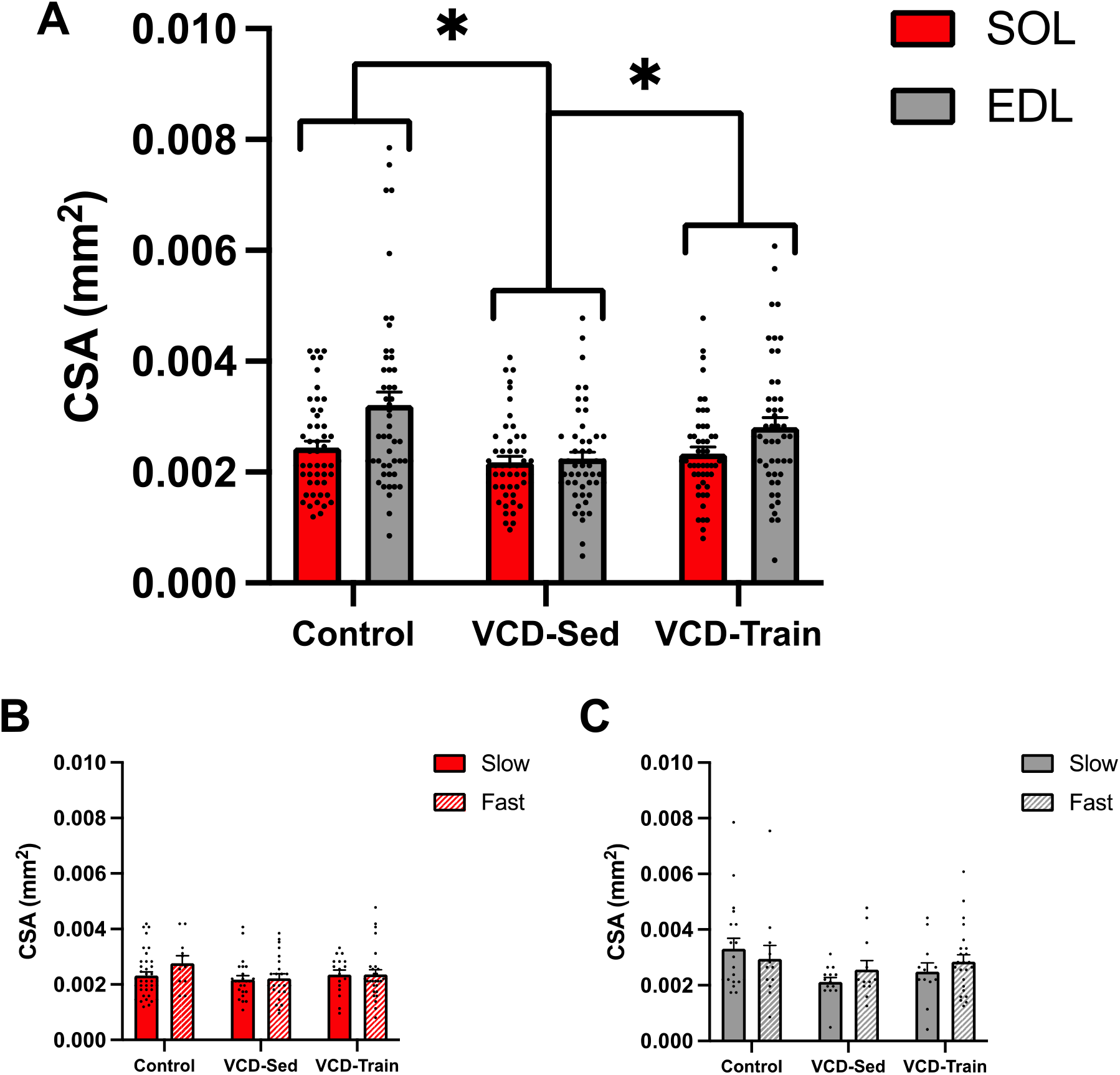
A) Cross-sectional area. Comparison of single SOL (red) and EDL (grey) muscle fibre CSA. VCD-sedentary fibres were smaller than control and VCD-training fibres and EDL fibres were bigger than SOL fibres. **B) SOL muscle.** When analyzing the SOL muscle by fibre phenotype, there were no differences across groups for either phenotype. **C) EDL muscle.** When analyzing the EDL muscle by fibre phenotype, there were no differences across groups for either phenotype. **\*** indicates significant difference (*p* < 0.05).

### Specific force

For force normalized to CSA, there was no interaction of group × muscle (*p* = 0.6256), but there was a main effect of group (*p* = 0.0148) such that VCD-sedentary fibres produced 11.6% less force than control fibres and 12.7% less force/CSA than VCD-training fibres (Figure 5A). There was also a main effect of muscle (*p* < 0.0001) such that EDL fibres produced 18.4% less force/CSA than SOL fibres. For the SOL, when fibres were binned based on V_max_, there was no interaction of group × phenotype (*p* = 0.6027), nor main effect of group (*p* = 0.1756) (Figure 5B). There was an effect of phenotype such that fast SOL fibres produced 20.3% more force/CSA compared to slow SOL fibres. For the EDL, when fibres were binned based on V_max_, there was no interaction of group × phenotype (*p* = 0.5755), nor main effects of group (*p* = 0.5029) or phenotype (*p* = 0.6830) (Figure 5C).

**Figure 5:**
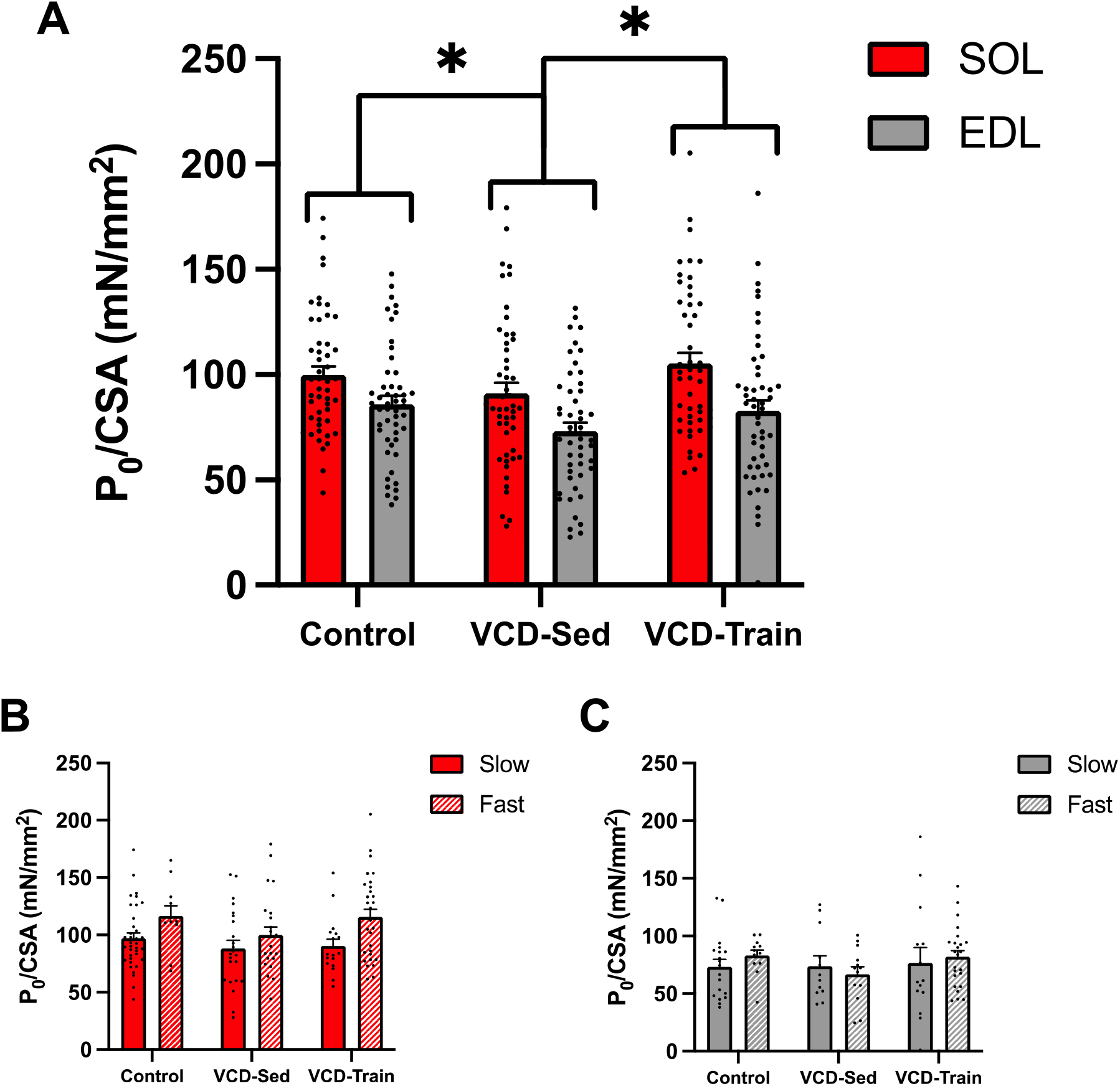
A) Specific force. Comparison of peak isometric force from maximally activated (pCa 4.5) single SOL (red) and EDL (grey) muscle fibres normalized to CSA (P_0_/CSA, mN/mm^2^). VCD-sedentary fibres produced less force/CSA compared to control and VCD-training fibres, and EDL fibres produced less force/CSA than SOL fibres. **B) SOL muscle.** When analyzing the SOL muscle by fibre phenotype, there were no differences across groups, but fast fibres (hashed) produced more force/CSA than slow fibres (solid). **C) EDL muscle.** When analyzing the EDL muscle by fibre phenotype, there were no differences across groups for either phenotype. **\*** indicates significant difference (*p* < 0.05).

### Maximum shortening velocity

For V_max_ from the fit force-velocity curve, there was an interaction of group × muscle (*p* = 0.0415) such that V_max_ was 25.0% slower in VCD-training EDL fibres compared to control EDL fibres (Figure 6A). There was no main effect of group (*p* = 0.2704), but there was a main effect of muscle (*p* = 0.0007) such that V_max_ was 27.3% faster in EDL fibres than SOL fibres. For the SOL, when fibres were binned based on V_max_, there was no interaction of group × phenotype (*p* = 0.8672), but there was an effect of group (*p* = 0.0015) such that V_max_ was 15.4% slower in VCD-sedentary fibres than control fibres, and 14.0% slower in VCD-training fibres than control fibres (Figure 6B). For the EDL, when fibres were binned based on V_max_, there was an interaction of group × phenotype (*p* = 0.0238) and a main effect of group (p = 0.0004) such that V_max_ was 27.2% slower in fast VCD-sedentary fibres compared to fast control fibres, and 43.6% slower in fast VCD-training fibres compared to fast control fibres (Figure 6C). As V_max_ was used to phenotypically bin fibres based on contractile speeds, there was the expected effect of phenotype in both SOL (*p* < 0.0001) and EDL (*p* < 0.0001) fibres with V_max_ being 82.9 and 136.7% faster in fast fibres than slow fibres for the SOL and EDL, respectively (Figure 6B&C).

**Figure 6:**
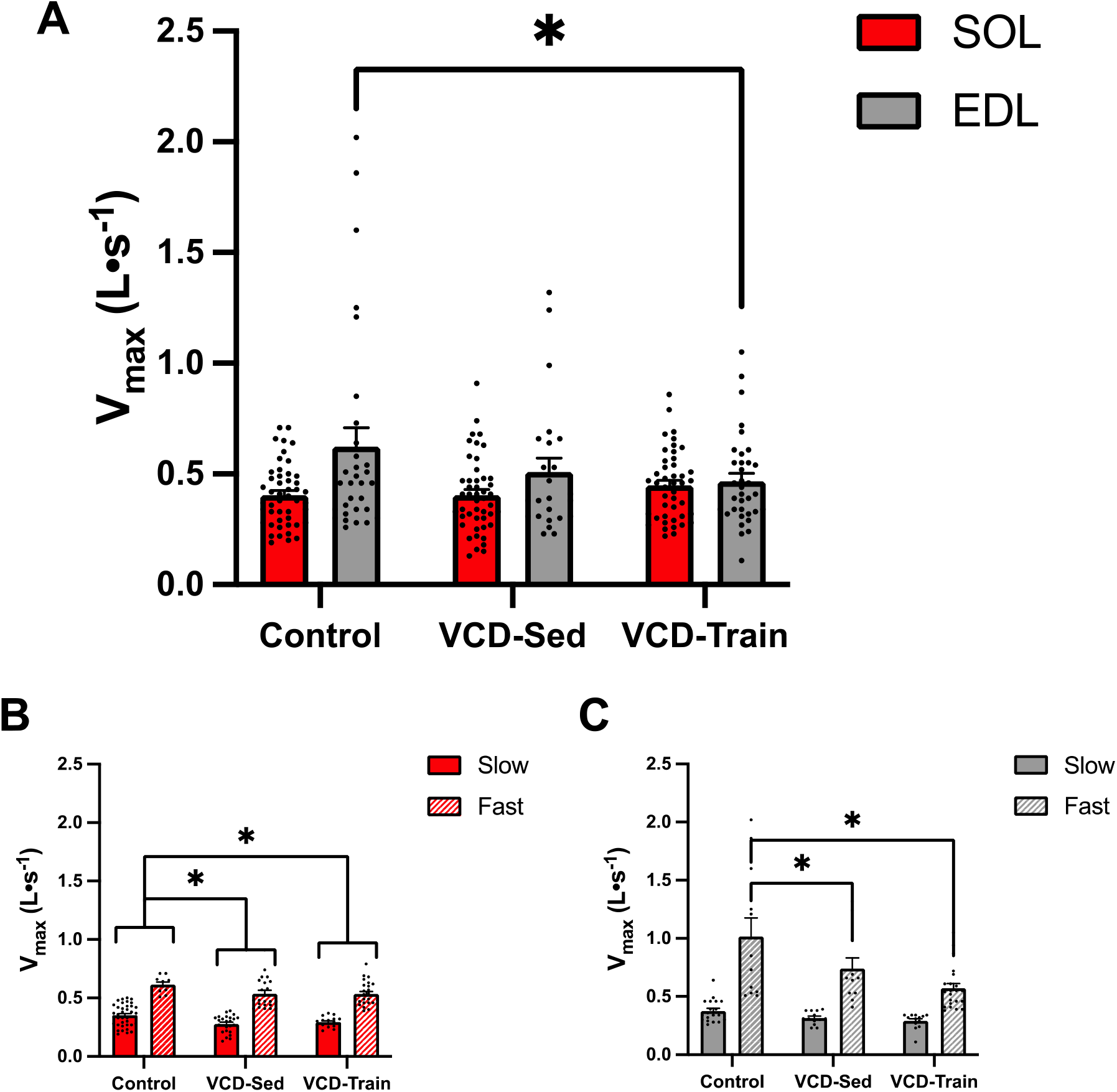
A) Maximum shortening velocity. Comparison of V_max_ (L_0_•s^-1^) determined from fit force-velocity-power curves from single SOL (red) and EDL (grey) muscle fibres. V_max_ was slower in VCD-training EDL fibres than control EDL fibres. **B) SOL muscle.** When analyzing the SOL muscle by fibre phenotype, VCD-sedentary and VCD-training fibres both had slower V_max_ values than control fibres. **C) EDL muscle.** When analyzing the EDL muscle by fibre phenotype, V_max_ was slower in fast VCD-sedentary fibres and fast VCD-training fibres compared to fast control fibres. **\*** indicates significant difference (*p* < 0.05).

### Absolute peak power

For peak power from the fit force-power curve, there was no interaction of group × muscle (*p* = 0.3030), but there was a main effect of group (*p* = 0.0004) such that peak power was 32.2% lower in VCD-sedentary fibres than in control fibres (Figure 7A). There was also a main effect of muscle (*p* = 0.0010) such that peak power was 28.5% higher in EDL fibres than in SOL fibres. For the SOL, when fibres were binned based on V_max_, there was an interaction of group × phenotype (*p* = 0.0170) and main effects of group (*p* < 0.0001) and phenotype (p < 0.0001) such that, in slow fibres, peak power was 27.2% lower in VCD-sedentary fibres than control fibres and, in fast fibres, peak power was 41.3% lower in VCD-sedentary fibres compared to controls, but 21.7% higher in VCD-training fibres compared to VCD-sedentary fibres. Peak power was also 28.6% lower in fast VCD-training fibres than fast control fibres (Figure 7B). For the EDL, when fibres were binned based on V_max_, there was no interaction of group × phenotype (*p* = 0.4624), but there was a main effect of group (*p* = 0.0090) such that peak power was 32.7% lower in VCD-sedentary EDL fibres compared to control EDL fibres and 29.2% lower in VCD-training EDL fibres compared to control EDL fibres (Figure 7C). There was also a main effect of phenotype (*p* = 0.0003) such that peak power was 55.9% higher in fast EDL fibres than slow EDL fibres.

**Figure 7:**
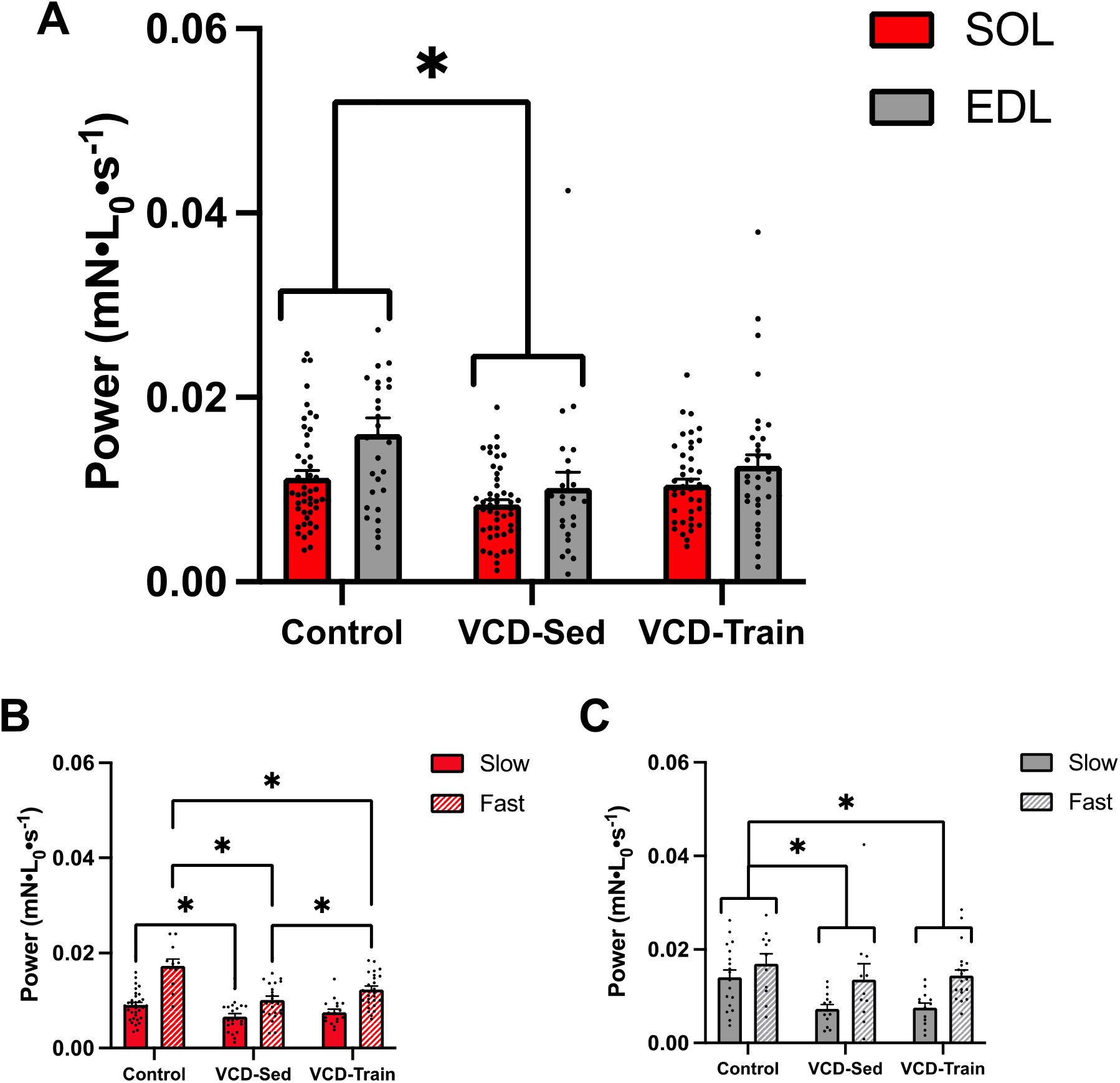
A) Peak power. Comparison of peak power (mN•L_0_•s^-1^) determined from the fit force-velocity-power curves from single SOL (red) and EDL (grey) muscle fibres. Fit peak power was lower in VCD-sedentary fibres compared to controls and EDL fibres had higher peak power than SOL fibres. **B) SOL muscle.** When analyzing the SOL muscle by fibre phenotype, slow (solid) VCD-sedentary fibres had lower peak power than slow controls, and fast (hashed) VCD-sedentary fibres had lower peak power than fast control and VCD-training fibres. Fast VCD-training fibres also had lower peak power than fast control fibres. **C) EDL muscle.** When analyzing the EDL muscle by fibre phenotype, VCD-sedentary and VCD-training fibres both had lower peak power compared to control fibres, and peak power was higher in fast fibres compared to slow fibres. **\*** indicates significant difference (*p* < 0.05).

### Specific peak power

For peak power from the force-power curve normalized to CSA, there was no interaction of group × muscle (*p* = 0.6148), nor was there an effect of group (*p* = 0.1749) (Figure 8A). There was, however, a main effect of muscle (*p* = 0.0363) such that peak power/CSA was 14.7% higher in EDL fibres than in SOL fibres. For the SOL, when fibres were binned based on V_max_, there was no interaction of group × phenotype (*p* = 0.7348), but there was a main effect of group (*p* = 0.0006) such that peak power was 27.2% lower in VCD-sedentary fibres than in control fibres and 17.6% lower in VCD-training fibres than in control fibres (Figure 8B). There was also an effect of phenotype (*p* < 0.0001) such that peak power was 50.2% higher in fast fibres compared to slow fibres. For the EDL, when fibres were binned based on V_max_, there was no interaction of group × phenotype (*p* = 0.9934), nor was there an effect of group (*p* = 0.5417) (Figure 8C). However, there was a main effect of phenotype (*p* < 0.0001) such that peak power was 66.2% higher in fast fibres compared to slow fibres.

**Figure 8:**
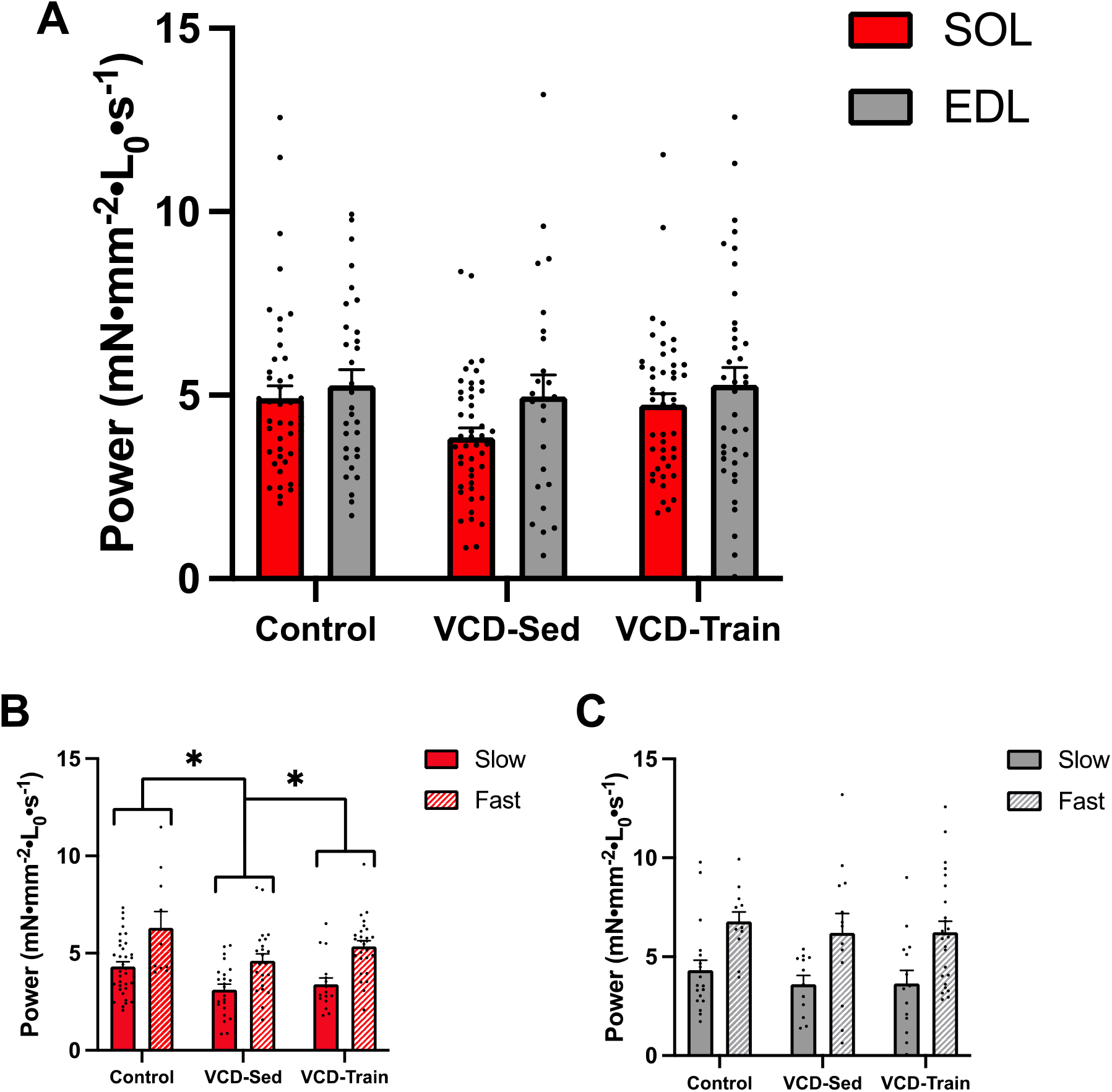
A) Specific peak power. Comparison of peak power determined from the fit force-velocity-power curves from single SOL (red) and EDL (grey) muscle fibres normalized to CSA (mN•mm^-2^•L_0_•s^-1^). There were no differences in specific peak power across groups for either muscle, but peak power/CSA was higher in EDL fibres than SOL fibres. **B) SOL muscle.** When analyzing the SOL muscle by fibre phenotype, VCD-sedentary and VCD-training fibres had lower peak power/CSA than control fibres, and fast fibres (hashed) had higher peak power/CSA than slow (solid) fibres. **C) EDL muscle.** When analyzing the EDL muscle by fibre phenotype, there were no differences in specific peak power across groups, but fast fibres had higher peak power/CSA than slow fibres. **\*** indicates significant difference (*p* < 0.05).

### Curvature

For curvature, determined as a/P_0_ from the Hill equation, there was no interaction of group × muscle (*p* = 0.2686), nor was there an effect of group (*p* = 0.2747) (Figure 9A), indicating that despite the reduction in absolute force, there was no shift to faster velocities in response to the lighter loads. There was, however, a main effect of muscle (*p* < 0.0001) such that curvature was 50.1% greater in EDL fibres than in SOL fibres. For the SOL, when fibres were binned based on V_max_, there was no interaction of group × phenotype (*p* = 0.3357), nor was there an effect of group (*p* = 0.9073) (Figure 9B). However, there was a main effect of phenotype (*p* < 0.0001) such that curvature was 49.9% lower in fast fibres compared to slow fibres. For the EDL, when fibres were binned based on V_max_, there was no interaction of group × phenotype (*p* = 0.7638), nor was there an effect of group (*p* = 0.1859) (Figure 9C). However, there was a main effect of phenotype (*p* < 0.0001) such that curvature was 42.1% lower in fast fibres compared to slow fibres.

**Figure 9:**
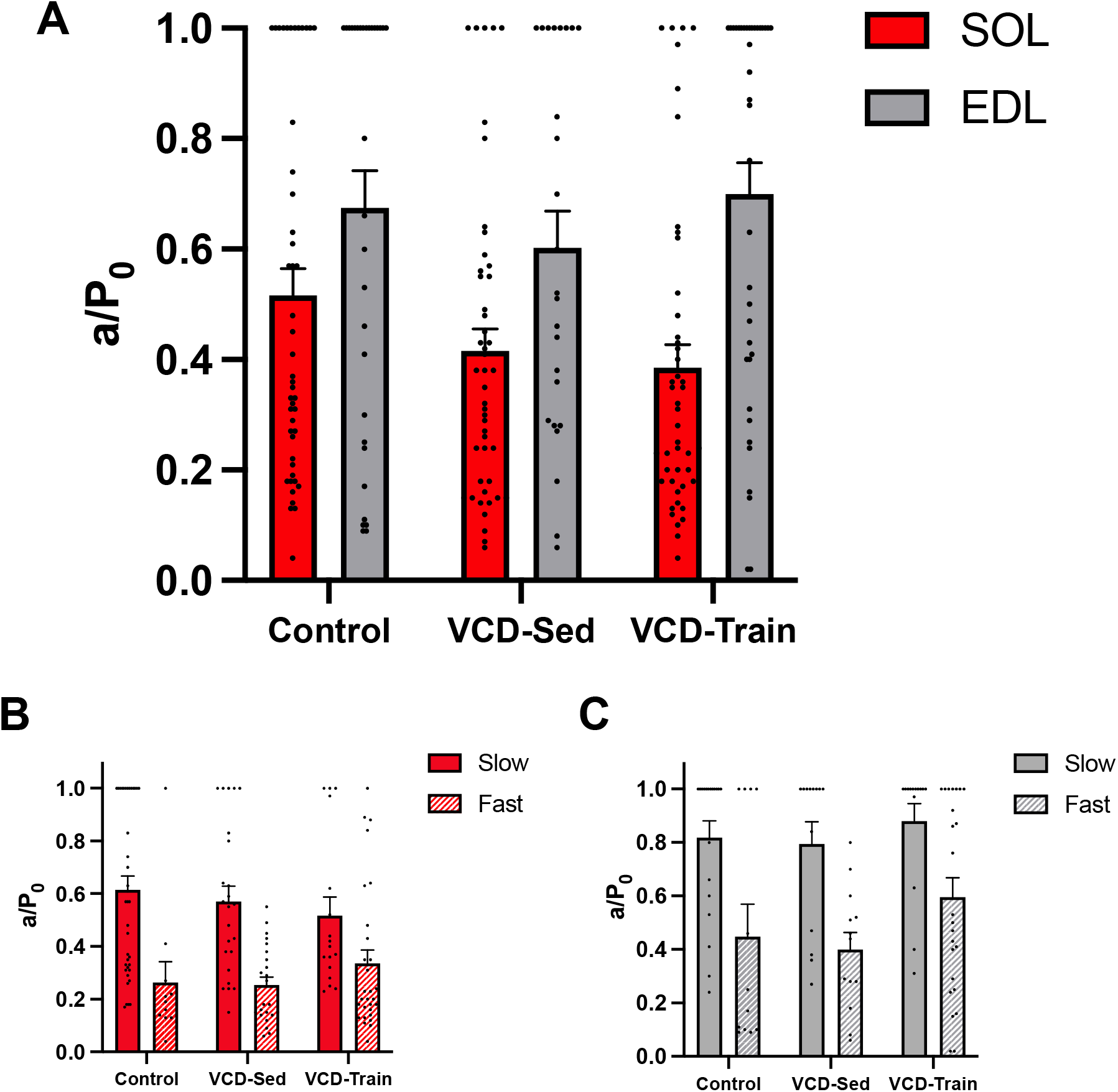
A) Curvature. Comparison of curvature (a/P_0_) from single SOL (red) and EDL (grey) muscle fibres. There were no differences in curvature across groups, but curvature was higher in EDL fibres compared to SOL fibres. **B) SOL muscle.** When analyzing the SOL muscle by fibre phenotype, there were no differences in curvature across groups, but curvature was lower in fast (hashed) fibres than slow (solid) fibres. **C) EDL muscle.** When analyzing the EDL muscle by fibre phenotype, there were no differences in curvature across groups, but curvature was lower in fast fibres than slow fibres. **\*** indicates significant difference (*p* < 0.05).

### Fibre phenotype distribution

For the SOL muscle, there was a shift to a higher proportion of fast fibres from the control group (22% fast fibres) to the VCD-sedentary (50% fast fibres) and the VCD-training groups (63% fast fibres) (Figure 10A). For the EDL muscle, there was a smaller change in fibre type distribution between the control (39% fast fibres) and VCD-sedentary group (46% fast fibres) with a greater proportion of fast fibres (64% fast fibres) in the VCD-training group (Figure 10B).

**Figure 10:**
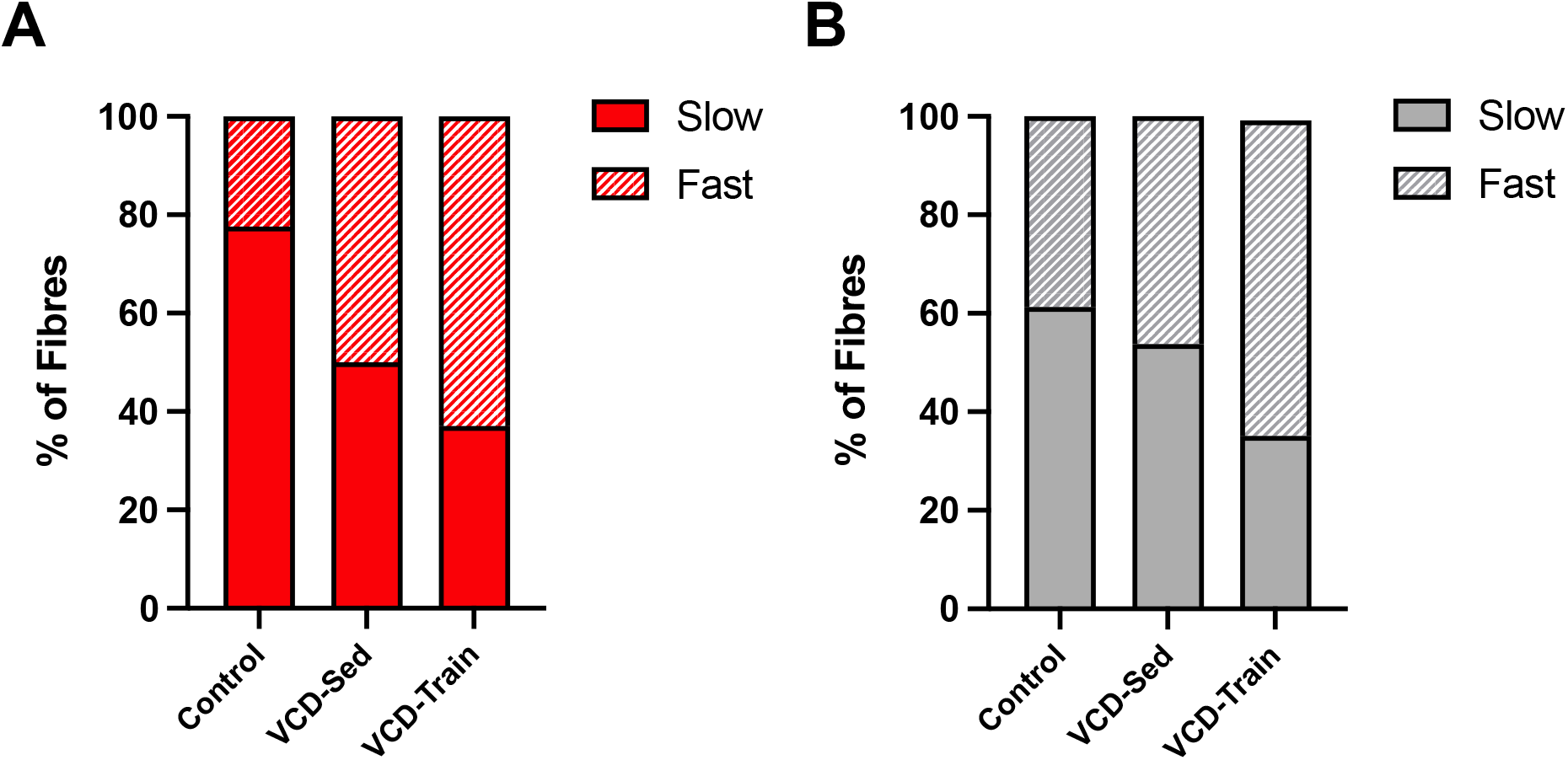
Fibre phenotype distribution. Comparison of fibre phenotype distribution (%) between groups. **A) SOL muscle.** For the SOL muscle, there was a shift to a higher proportion of fast fibres from the control group (22% fast fibres) to the VCD-sedentary (50% fast fibres) and the VCD-training groups (63% fast fibres). **B) EDL muscle.** For the EDL muscle, there was a smaller change in fibre type distribution between the control (39% fast fibres) and VCD-sedentary group (46% fast fibres) with a greater proportion of fast fibres (64% fast fibres) in the VCD-training group.

## Discussion

In the present study, we investigated the effects of gradual ovarian failure on muscle contractility, with an emphasis on power production, and the ability of HIIT to mitigate impairments. We found that VCD-induced ovarian failure reduced absolute peak power by ∼32.2% across muscles and that HIIT was insufficient to reverse this reduction, with the exception of fast SOL fibres where training partially attenuated (17.8%) absolute peak power loss. When peak power was normalized to CSA, VCD-induced impairments were only present in SOL muscles once assessed by mechanical phenotype. Our VCD model of menopause allowed us to evaluate the effects of ovarian failure on single muscle fibre function without the removal of the ovaries or halting ovarian androgen production, thereby more closely recapitulating the human condition, and providing novel insight about menopause-induced impairments to dynamic muscle function and possible tools to mitigate these deleterious effects.

### Ovarian failure impaired static components of single muscle fibre contractile function

Following ovarian failure, we observed a 31.0% reduction in single muscle fibre absolute force (Figure 3) as compared with controls. This finding is in agreement with some (6, 7, 15), but not all previous work assessing the effects of premature ovarian insufficiency on muscle contractile function in rodents (5, 9, 33, 34). Moran et al. (7) observed an 18% reduction in maximally activated force in intact SOL and EDL muscles from mature OVX mice (6 months of age, 8-9 weeks post-surgery), along with a 25% reduction in EDL fibre absolute force. Similarly, at 10 and 14 weeks after OVX, Wattanapermpool & Reiser (6) reported a 19 and 20% decrease in absolute force generated by SOL single muscle fibres, respectively. Using a skeletal muscle estrogen receptor knockout model of ovarian hormone reduction, Cabelka et al. (15) found that isometric, concentric and eccentric torque were all 7-8% lower in the plantar flexors of knockout mice as compared to wild type mice. Like in the present study, these findings from other models of ovarian failure show impairments in static contractile function at the levels of both single fibres and intact muscles owing to a reduction in circulating hormone levels. In a recent study from our lab, Mashouri et al. (5) found that VCD-induced ovarian failure increased absolute force of single muscle fibres from the SOL by 33%, but this study tested fibres immediately following the peri-menopausal transition, representing the onset of ovarian failure, at which point the effects of chronic ovarian hormone deficiency would not have been observable. Moreover, Greising et al. (9) found no differences in intact SOL muscle absolute force between control and VCD-treated mice 12 weeks after VCD injection, representing a significantly later menopause status than the one achieved by the current study. Therefore, it appears that the time after ovarian failure may explain at least some differences in findings and that impairment at the single muscle fibre level may not always translate to impairment in intact muscle. Future studies should seek to investigate time-course changes to accurately assess the impact of time as a variable.

When normalized to fibre CSA, we found a smaller albeit still significant reduction in specific force in the VCD-sedentary mice as compared with controls (Figure 5), corresponding to the smaller CSAs observed in VCD-sedentary fibres as compared with controls (Figure 4). This decrease in fibre CSA does not agree with previous studies which report either no effect of ovarian hormone status on muscle fibre CSA (35), or an increase in fibre CSA following OVX (36). This divergence from the literature suggests that some of the effect of VCD-induced ovarian failure on contractile function observed in the present study may have been owing to a decrease in fibre size and therefore a reduction in the number of available cross-bridges. Previous work from our lab observed increased absolute force production in the SOL muscle at the onset of ovarian failure; however, no differences between groups were present when force was normalized to CSA, indicating that it is unlikely that Ca^2+^ regulation of force is driving changes in force-generating capacity (5). Rather, it is possible that an initial compensatory hypertrophy occurs at the onset of menopause to offset functional impairments, followed by gradual muscle atrophy resulting in an overall decrease in fibre size and strength.

### Ovarian failure impaired dynamic components of single muscle fibre contractile function

Following ovarian failure, we observed a 32.2% reduction in peak power as compared with controls (Figure 7); however, peak power normalized to CSA did not differ between groups (Figure 8). Conversely, when fibres from the SOL muscle were assessed based on phenotype (i.e., contractile speed), there was still a reduction in peak power/CSA in both trained and sedentary VCD-treated mice (Figure 8B). Accordingly, the impairment in absolute dynamic function was most prominent in fast SOL fibres (Figure 7B). These findings differ from previous work using other models of ovarian failure. In *in vitro* studies of SOL and EDL whole muscles from OVX mice, no changes in peak power were observed (7). However, Moran et al. (7) also reported a decrease in EDL single fibre force without a concomitant increase in contractile velocity in the EDL muscle, so it is possible that power may have been impaired at the single fibre level. SOL muscle power was preserved owing to an increased shortening velocity following OVX (7), an adaptation that was not seen in the present study (Figure 6).

In another model of ovarian failure, the absence of skeletal muscle estrogen receptors resulted in a 11-12% decrease in power during slow shortening velocities in mouse plantar flexors *in vivo*, without any effect on peak power, thereby suggesting that estrogen deficiency-induced functional impairment was likely driven by a reduction in force generating capacity rather than a decrease in velocity (15). In contrast, the present study revealed significant effects of VCD-induced ovarian failure on peak power (Figure 7), and power at all loads between 20 and 70% of P_0_ (Supplementary Figures S1&S2), indicating that there may indeed have been impairments in the velocity component as well. Accordingly, given the lower absolute forces produced by the VCD-sedentary fibres, higher contractile velocities would have been expected owing to the hyperbolic force-velocity relationship of muscle (27, 37). However, despite the lower absolute loads, there was no change in curvature (Figure 9) nor any difference in shortening velocities between groups across loads (Supplementary Figures S3&S4). Moreover, V_max_ was impaired in the VCD-sedentary and VCD-training groups (Figure 6). As such, we can elucidate that distinct effects of ovarian failure may depend on either the presence or absence of ovarian tissue, or the level (cellular or whole muscle) at which impairments are assessed.

### HIIT restored force production, but not peak power

Despite ovarian failure, reductions in both absolute and normalized force were attenuated following HIIT such that absolute force was not significantly different between control and VCD-training fibres (Figures 3&5). This finding is consistent with previous training studies in models of premature ovarian insufficiency and/or aging. Greising et al. (9) found an effect of running in aged ovarian-senescent mice such that *in vitro* SOL force production and active stiffness improved following 8 weeks of voluntary wheel running in hormone-replaced and non-replaced mice. Similarly, 16 weeks of voluntary wheel running increased CSA, peak twitch force, and passive stiffness in *in vitro* SOL muscles from skeletal muscle estrogen receptor-ablated mice (15). In an investigation of the ability of HIIT to offset sarcopenia in aged male and female mice, a training protocol closely resembling the one used in the present study successfully increased muscle mass and single fibre CSA, and improved grip force, treadmill endurance, sprint performance, and gait speed (19, 20).

Although muscle power was not assessed directly in the above cited HIIT studies, their findings indicate that improvements in both the force and velocity components were present, which would point to an increase in power as assessed by performance metrics (19). In the present study, there was no attenuation of peak power loss with training, except in fast SOL fibres where peak power loss was partially attenuated, but not to the level of controls (Figure 7). Interestingly, fast-type fibres from the SOL also had the most power loss following ovarian failure, meaning that the most impaired fibres benefitted the most from the HIIT intervention. It remains unclear whether this muscle and type-specific attenuation of impairment was simply due to a greater influence of ovarian failure on fast SOL fibres, and therefore more room for functional improvement with training, or rather owing to the specific training paradigm stimulating adaptations in these fibres alone. Given the lack of response to HIIT in the EDL muscle, and the slower V_max_ observed in both VCD-sedentary and VCD-training fibres (Figure 6), we can conclude that our exercise paradigm had a predominantly endurance-based effect which would have provided a strong stimulus for adaptation of SOL size and strength, but not for increased velocity in either muscle. A greater intensity of training or the addition of resistance while running could help ascertain whether HIIT is capable of recovering velocity following ovarian failure. Furthermore, the HIIT program described by Seldeen et al. (23) was twice as long as the one employed here, so it is possible that the length of our intervention was simply insufficient to modulate dynamic contractile function. Alternatively, the effects of HIIT may not have been observable at the single muscle fibre level. In a comparison of lifelong exercisers against age-matched non-exercisers, Gries et al. (17) found that single fibre power was significantly higher in those who had been exercising regularly for ∼50 years, but that these adaptations did not appear to translate to whole muscle size or function. As such, it seems that there may be distinct mechanisms at play at the single fibre level as compared to the joint level.

### Mechanical Phenotypic shift and fibre type-specific adaptations as mechanisms of maintaining power production

We observed an increase in the proportion of fast-type SOL fibres following VCD injection and a further shift to fast-type fibres in the SOL muscle with HIIT (Figure 10A). In the EDL muscle, on the other hand, there was minimal change in single muscle fibre mechanical phenotype distribution in the VCD-sedentary group, but an increase in the proportion of fast EDL fibres following training (Figure 10B). This shift in mechanical phenotype may explain why the magnitude of ovarian failure-induced impairment was lower in SOL fibres compared to EDL fibres, and why attenuation of power loss with HIIT was exclusively present in fast SOL fibres. Shifts in fibre type, particularly in the SOL muscle, have previously been reported in VCD-treated animals (3, 5), but not in OVX models which instead have shown either no change in SOL MHC isoform composition (33) or a reduction in fast MHC expression in the SOL muscle (38). Much like the present study, Perez et al. (3) observed a reduction in MHC type I fibres and an increase in MHC type II fibres in mouse SOL muscle following VCD treatment, along with an additional shift to faster MHC composition with voluntary wheel running. This adaptation was specific to the SOL muscle as no changes in MHC isoform composition were observed in the tibialis anterior, gastrocnemius, or quadriceps muscles (3). Mashouri et al. (5) suggested that a shift to a greater proportion of MHC type II fibres in the SOL muscle was likely driving the increase in absolute force observed at the onset of chemically induced ovarian failure and suggested that the SOL muscle was quicker to respond to the reduction of circulating 17β-estradiol. However, given that the same phenotypic shift in the SOL and lack thereof in the EDL occurred in the present study an additional 8 weeks after ovarian failure (i.e., late menopause), it appears that VCD treatment may just not trigger changes in the fast-type EDL muscle MHC composition or phenotypical behaviour.

When assessing independent measures of force and velocity, Gries et al. (17) observed that lifelong exercisers exhibited greater MHC type I fibre force than non-exercisers with no difference in contractile velocity, and lower MHC type II fibre force compared to young healthy individuals with an increase in maximum contractile velocity. Based on these fibre-type specific findings, it has been suggested that the exercise-induced preservation of single fibre power may be driven by improved force production in MHC type I fibres, and increased contractile velocity in MHC type II fibres (17). In the present study, the same improvement in force production occurred following training, which likely contributed to the partial attenuation of power loss in SOL muscle fibres. Contrarily, there was no effect of ovarian failure or training to improve contractile velocity across loads (Supplementary Figure S3&S4) – apart from SOL fibres at 10 and 20% of L_0_ (Supplementary Figure S3) – which may explain the failure of HIIT to completely attenuate impairments in dynamic single fibre function.

## Conclusion

Using a VCD model of ovarian failure to better approximate the natural progression of menopause, we found that ovarian failure impairs dynamic contractile function at the single muscle fibre level. Absolute force and peak power were reduced by 31.0 & 32.2%, respectively, in VCD-sedentary fibres as compared to controls across muscles and fibre phenotypes. The observed impairment in single muscle fibre power appears to be driven by some combination of a reduction of force-generating capacity, owing to a reduction in fibre size, and a blunted shift to faster shortening velocities. Our uphill HIIT program completely offset impairments in force production, but only partially attenuated power impairments. Nonetheless, exercise may still be a useful tool to counteract some of the deleterious effects on skeletal muscle associated with menopause.

## Acknowledgements

We would like to thank Benjamin Dalton and Avery Hinks for their help with data processing, and Matt Borkowski for technical assistance with establishing the isotonic load clamp procedures.

## Ethics statement

All procedures were approved by the Animal Care Committee (AUP 4714) of the University of Guelph.

## Data accessibility

Individual values of all supporting data are available upon request.

## Author contributions

All authors contributed equally.

---

**Supplementary Figure S1:**
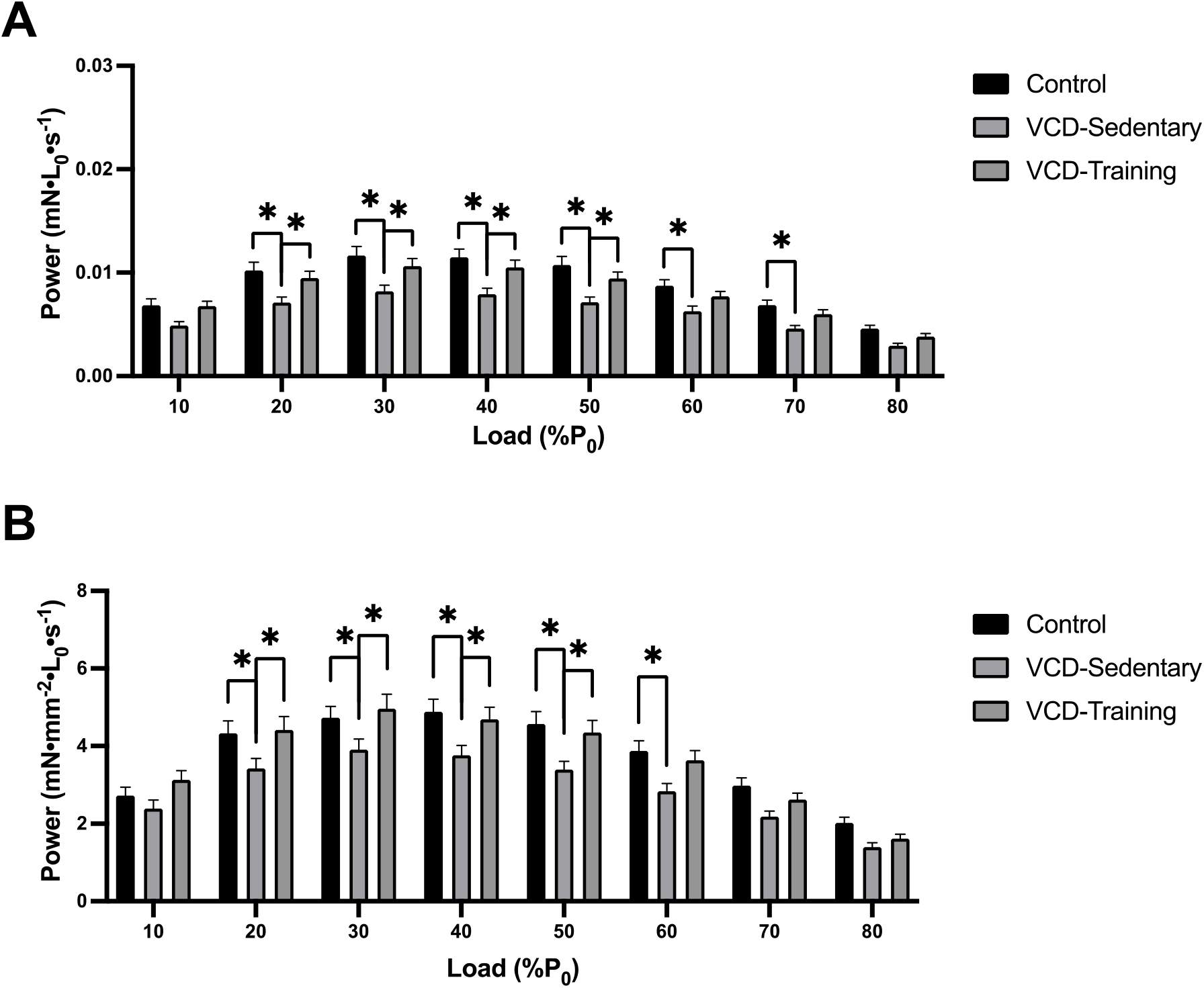
A) SOL power across loads. Comparison of actual experimental single SOL muscle fibre power (mN•L_0_•s^-1^) achieved during clamped loads (10-80% of P_0_). Power was lower in VCD-sedentary fibres compared to control and VCD-training fibres for all loads between 20 and 70% of P_0_. **B) Specific SOL power across loads.** When actual experimental single SOL muscle fibre power was normalized to CSA (mN•mm^-2^•L_0_•s^-1^), power remained lower in VCD-sedentary fibres compared to control and VCD-training fibres for all loads between 20 and 50% of P_0_, and was lower in VCD-sedentary fibres compared to control fibres only at 60% of P_0_. **\*** indicates significant difference (*p* < 0.05).

**Supplementary Figure S2:**
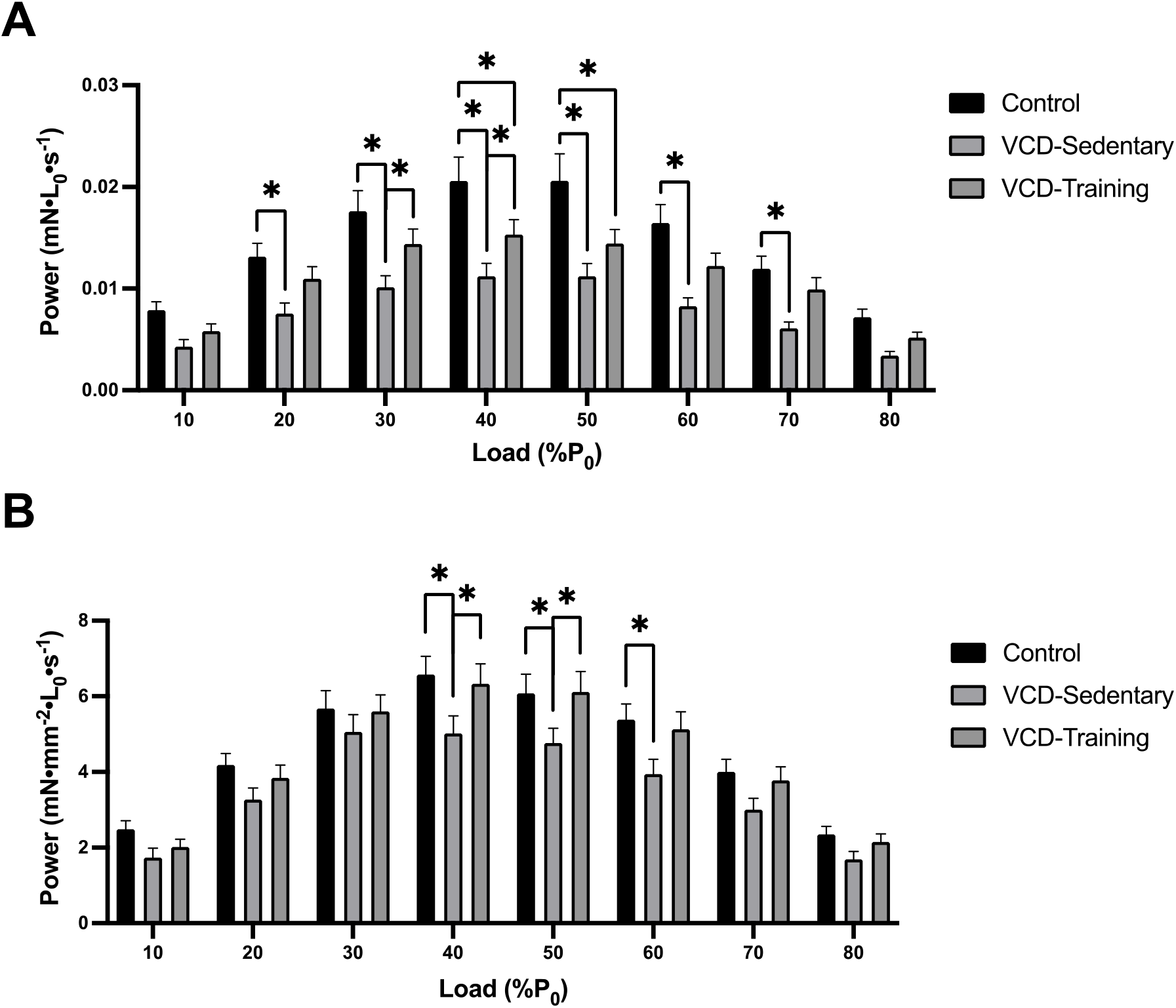
A) EDL power across loads. Comparison of actual experimental single EDL muscle fibre power (mN•L_0_•s^-1^) achieved during clamped loads (10-80% of P_0_). Power was lower in VCD-sedentary fibres compared to control fibres for all loads between 20 and 70% of P_0_, and in VCD-sedentary fibres compared to VCD-training fibres at 30 and 40% of P_0_. Power was also lower in VCD-training fibres compared to control fibres at 40 and 50% of P_0_. **B) Specific EDL power across loads.** When actual experimental single EDL muscle fibre power was normalized to CSA, power remained lower in VCD-sedentary fibres compared to control and VCD-training fibres at 40 and 50% of P_0_, and was lower in VCD-sedentary fibres compared to control fibres only at 60% of P_0_. **\*** indicates significant difference (*p* < 0.05).

**Supplementary Figure S3:**
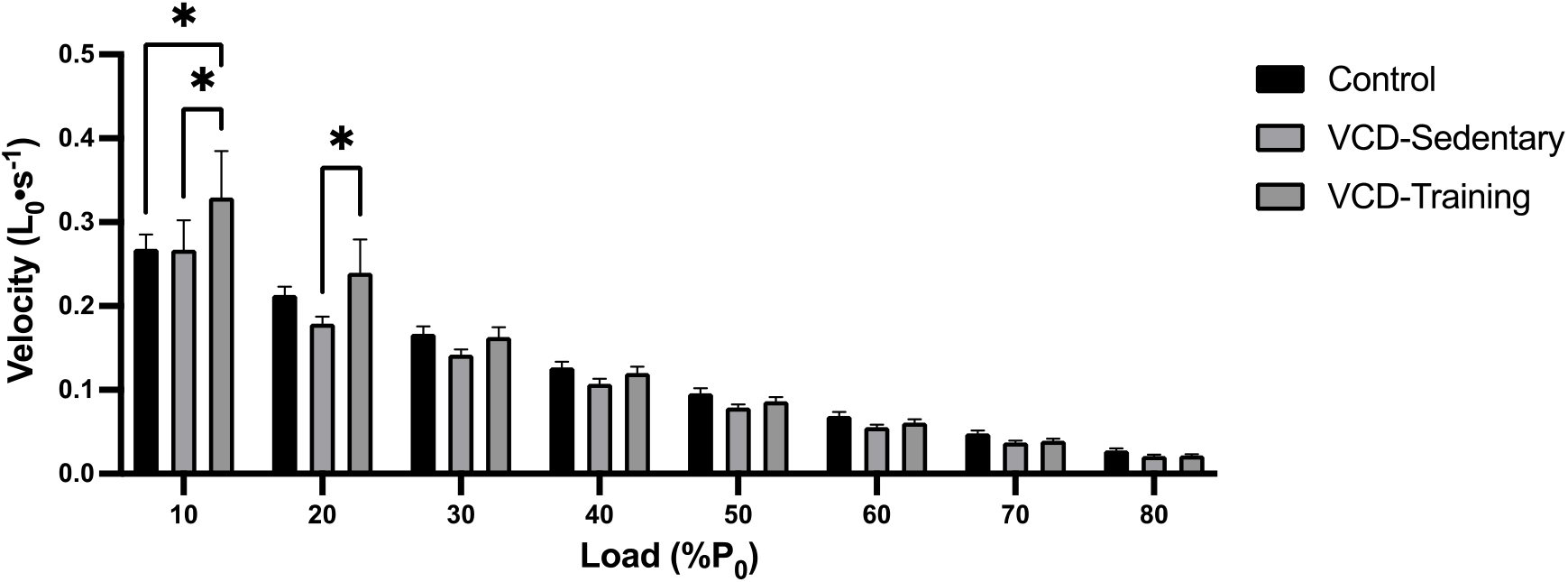
SOL velocity across loads. Comparison of actual experimental single SOL muscle fibre contractile shortening velocity (L_0_•s^-1^) achieved during clamped loads (10-80% of P_0_). Shortening velocity was faster in VCD-training fibres compared to control and VCD-sedentary fibres at 10% of P_0_, and was faster in VCD-training fibres compared to VCD-sedentary fibres only at 20% P_0_. There were no differences at any other loads **\*** indicates significant difference (*p* < 0.05).

**Supplementary Figure S4:**
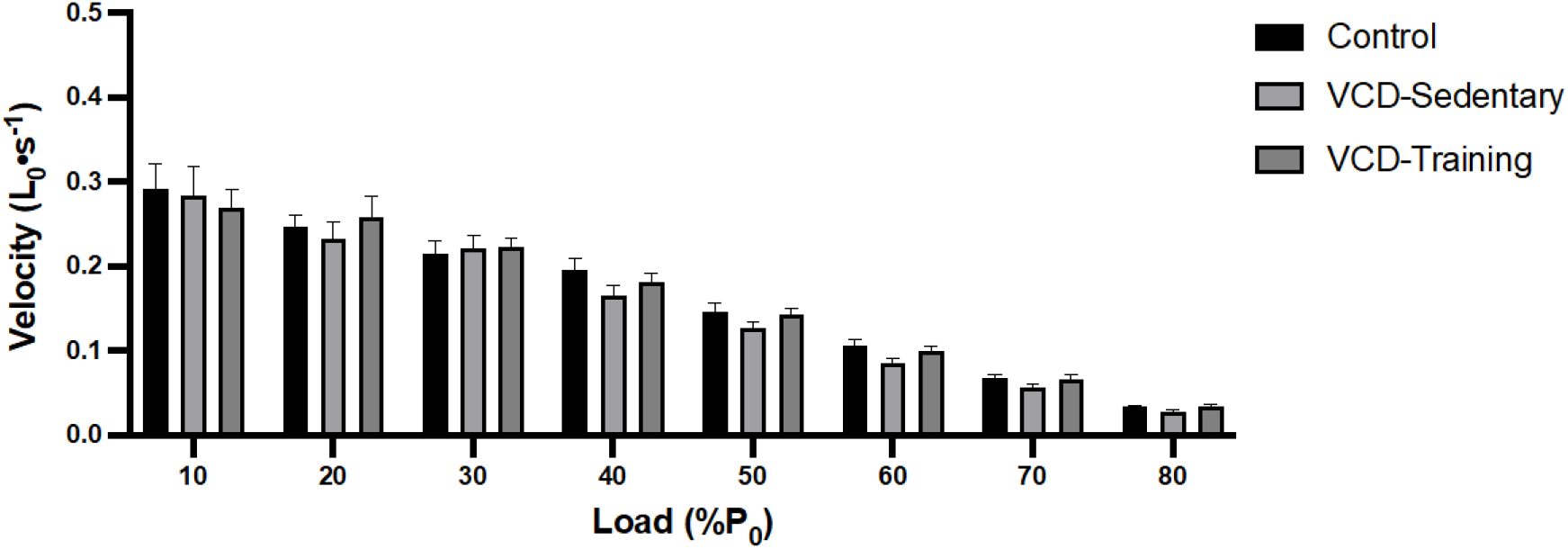
EDL velocity across loads. Comparison of actual experimental single EDL muscle fibre contractile shortening velocity (L_0_•s^-1^) achieved during clamped loads (10-80% of P_0_). There were no differences in shortening velocity between groups at any load. **\*** indicates significant difference (*p* < 0.05).

